# Development of a First-in-Class RIPK1 Degrader to Enhance Antitumor Immunity

**DOI:** 10.1101/2024.03.25.586133

**Authors:** Xin Yu, Dong Lu, Xiaoli Qi, Hanfeng Lin, Bryan L. Holloman, Feng Jin, Longyong Xu, Lang Ding, Weiyi Peng, Meng C. Wang, Xi Chen, Jin Wang

**Affiliations:** The Verna and Marrs McLean Department of Biochemistry and Molecular Pharmacology, Baylor College of Medicine, Houston, Texas 77030, USA; Department of Molecular and Cellular Biology, Baylor College of Medicine, Houston, Texas 77030, USA; Howard Hughes Medical Institute, Janelia Research Campus, Ashburn, Virginia 20147, USA; Department of Biology and Biochemistry, University of Houston, Houston, Texas 77204, USA

## Abstract

The scaffolding function of receptor interacting protein kinase 1 (RIPK1) confers intrinsic and extrinsic resistance to immune checkpoint blockades (ICBs) and has emerged as a promising target for improving cancer immunotherapies. To address the challenge posed by a poorly defined binding pocket within the intermediate domain, we harnessed proteolysis targeting chimera (PROTAC) technology to develop a first-in-class RIPK1 degrader, LD4172. LD4172 exhibited potent and selective RIPK1 degradation both *in vitro* and *in vivo*. Degradation of RIPK1 by LD4172 triggered immunogenic cell death (ICD) and enriched tumor-infiltrating lymphocytes and substantially sensitized the tumors to anti-PD1 therapy. This work reports the first RIPK1 degrader that serves as a chemical probe for investigating the scaffolding functions of RIPK1 and as a potential therapeutic agent to enhance tumor responses to immune checkpoint blockade therapy.

## Introduction

Immune checkpoint blockades (ICBs) have transformed cancer therapy by disrupting inhibitory signals that typically weaken robust anti-tumor immune responses ^1^. Despite the success of ICBs, a significant subset of patients remain unresponsive to ICBs owing to various immuno-resistances, which are often propagated by cancer cells ^2^. The exploration of combinational therapies involving novel immunomodulatory agents with anti-PD-1/PD-L1 has emerged as a promising approach to overcome intrinsic or acquired resistance to ICBs ^3^.

Receptor-interacting protein kinase 1 (RIPK1) regulates cell fate through its kinase-dependent and--independent functions and controls proinflammatory responses downstream of multiple innate immune pathways, including those initiated by tumor necrosis factor-α (TNF-α), toll-like receptor (TLR) ligands, and interferons (IFNs) ^4^. Recent studies have shown that genetic knockout of RIPK1 in cancer cells significantly sensitizes tumors to anti-PD1, leading to drastic changes in the tumor microenvironment, including increased infiltration of effector T cells, reduction of immunosuppressive myeloid cells, and enhanced immunostimulatory cytokine secretion ^5–7^. Notably, RIPK1-mediated ICB resistance requires ubiquitin scaffolding function through its intermediate domain instead of its kinase function. Genetic depletion of RIPK1, but not inactivation of its kinase domain, sensitizes B16F10 tumors to ICBs ^5,6^. Hence, targeting RIPK1 scaffolding functions holds promise as a strategy to synergize with ICBs to promote antitumor immunity.

While all RIPK1 inhibitors developed thus far have focused on inhibiting kinase function for the treatment of autoimmune, inflammatory, and neurodegenerative diseases ^8^, the development of inhibitors specifically targeting the intermediate domain of RIPK1 remains challenging due to the absence of a well-defined binding pocket within this domain. A proteolysis-targeting chimera (PROTAC) is a heterobifunctional molecule that binds both a targeted protein and an E3 ubiquitin ligase to facilitate the formation of a ternary complex, leading to ubiquitination and ultimate degradation of the target protein ^9^. Using PROTAC technology, we developed LD4172, a first-in-class highly potent and specific RIPK1 degrader, to abolish the scaffolding functions of RIPK1. We showed that LD4172 potently induces RIPK1 degradation with high specificity and substantially sensitizes multiple preclinical cancer models to anti-PD1 therapy.

## Results

### Development of RIPK1 Degrader LD4172

To develop RIPK1 PROTACs, we tested two types of RIPK1 binders: type II RIPK1 inhibitor **1** (also referred as T2I), which targets both the ATP-binding pocket and the allosteric hydrophobic back pocket ^10^, and type III RIPK1 inhibitor **2**, which only binds the hydrophobic back pocket of the kinase domain ^11^. To identify the ideal attachment sites for PROTAC linkers, we performed molecular docking of **1** with RIPK1, which revealed a solvent-exposed ethyl group in the 7H-pyrrolo[2,3-d] pyrimidine ring (**Fig. 1A**). The co-crystal structure of RIPK1 in complex with **2** (PDB: 6R5F) showed that the oxadiazole moiety in **2** was solvent-exposed, providing an ideal exit vector for linker attachment (**Fig. 1B**).

**Figure 1.**
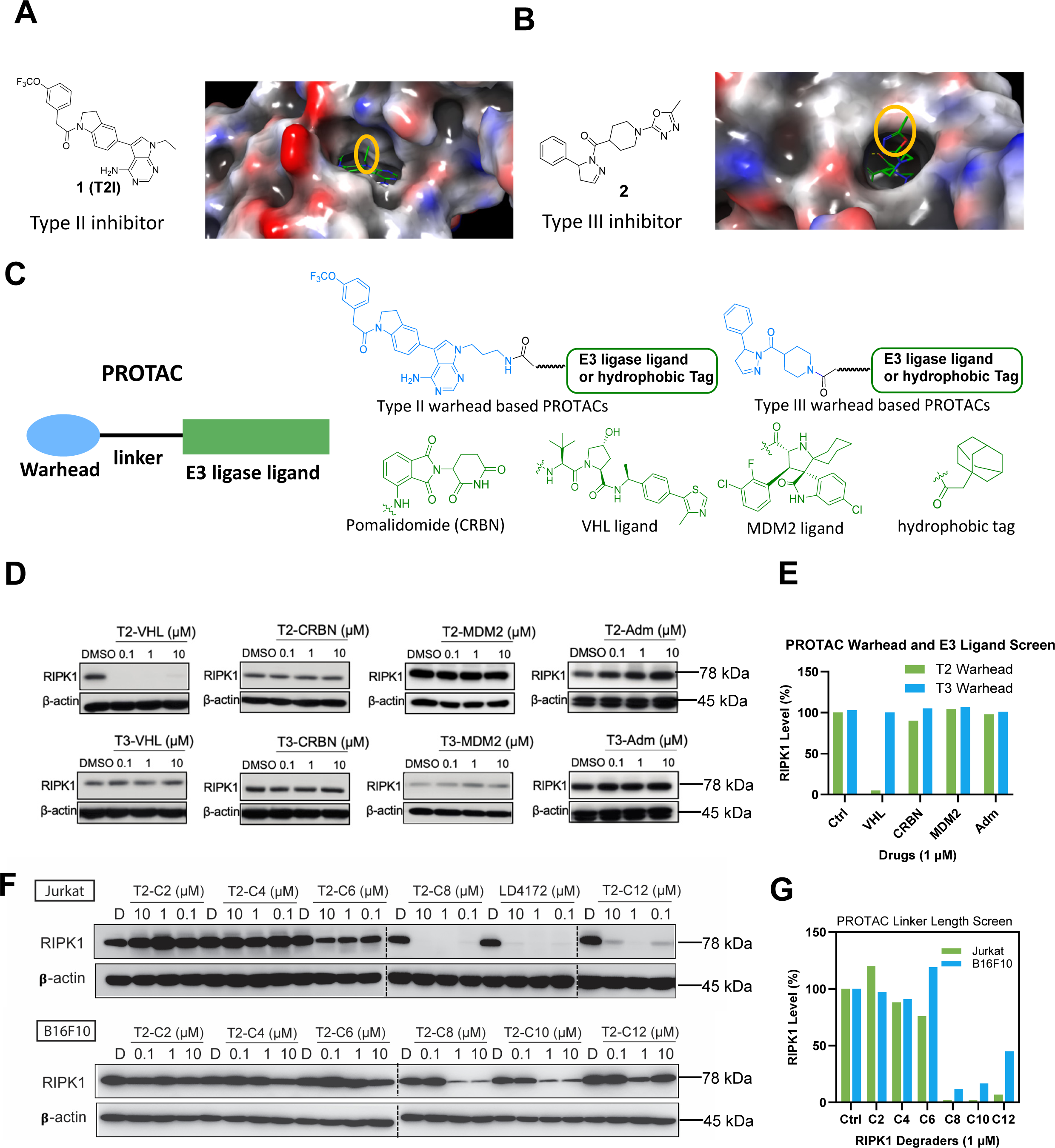
Design and screen of RIPK1 PROTACs. **A**, Chemical structure and docking modeling of type II inhibitor 1 with the kinase domain of RIPK1 (PDB: 4NEU). The group exposed to solvent region is highlighted with yellow circle. **B**, Chemical structure and co-crystal structure of type III inhibitor 2 in complex with RIPK1 (PDB: 6R5F). The solvent-exposed group of inhibitor 2 is highlighted with yellow circle. **C**, A small library design of RIPK1 PROTACs. Either Type II inhibitor 1 or type III inhibitor 2 is conjugated with various E3 ligase ligands or adamantane tag to generate a small library of RIPK1 PROTACs. **D**, Quantification of RIPK1 levels in Jurkat cells treated with indicated compounds for 24 h at 0, 0.1, 1 and 10 µM, followed by Western blotting. **E**, Quantification of **D**. **F**, Quantification of RIPK1 levels in both Jurkat and B16F10 cells treated with type II inhibitor-based PROTACs with different linker lengths for 24 h, followed by Western blotting. **G**, Quantification of **F**.

To identify an appropriate E3 ligase pair for RIPK1 degradation, we synthesized a small library by conjugating RIPK1 binders **1** and **2** to ligands for different E3 ligases, including Cereblon (CRBN), von Hippel-Lindau tumor suppressor (VHL), murine double minute 2 (MDM2), and a hydrophobic adamantane tag (**Fig. 1C**). As shown in **Fig.1D-E**, PROTACs formed by conjugating type II inhibitor **1** to a VHL ligand induced the most efficient degradation of RIPK1 in Jurkat cells.

We further optimized RIPK1 PROTACs through linker lengths ranging from two to 14 methylene groups (**Fig. 1F**). We found that PROTACs with linker lengths of more than six methylenes were able to effectively degrade >90% of RIPK1 at 1 µM after 24 h incubation in Jurkat cells, showing a monotonic trend (**Fig. 1F-G**). Consistently, the PROTACs exhibited maximal degradation with an 8-to 10-methylene linker and significantly reduced potency with either shorter or longer linkers in B16F10 mouse melanoma cells (**Fig. 1F-G**). Considering the potency of both human and mouse cells, we chose a combination of a type II RIPK1 binder, a VHL ligand, and a 10-methylene linker as the lead RIPK1 degrader, designated as LD4172 (**Fig. 2A**).

**Figure 2.**
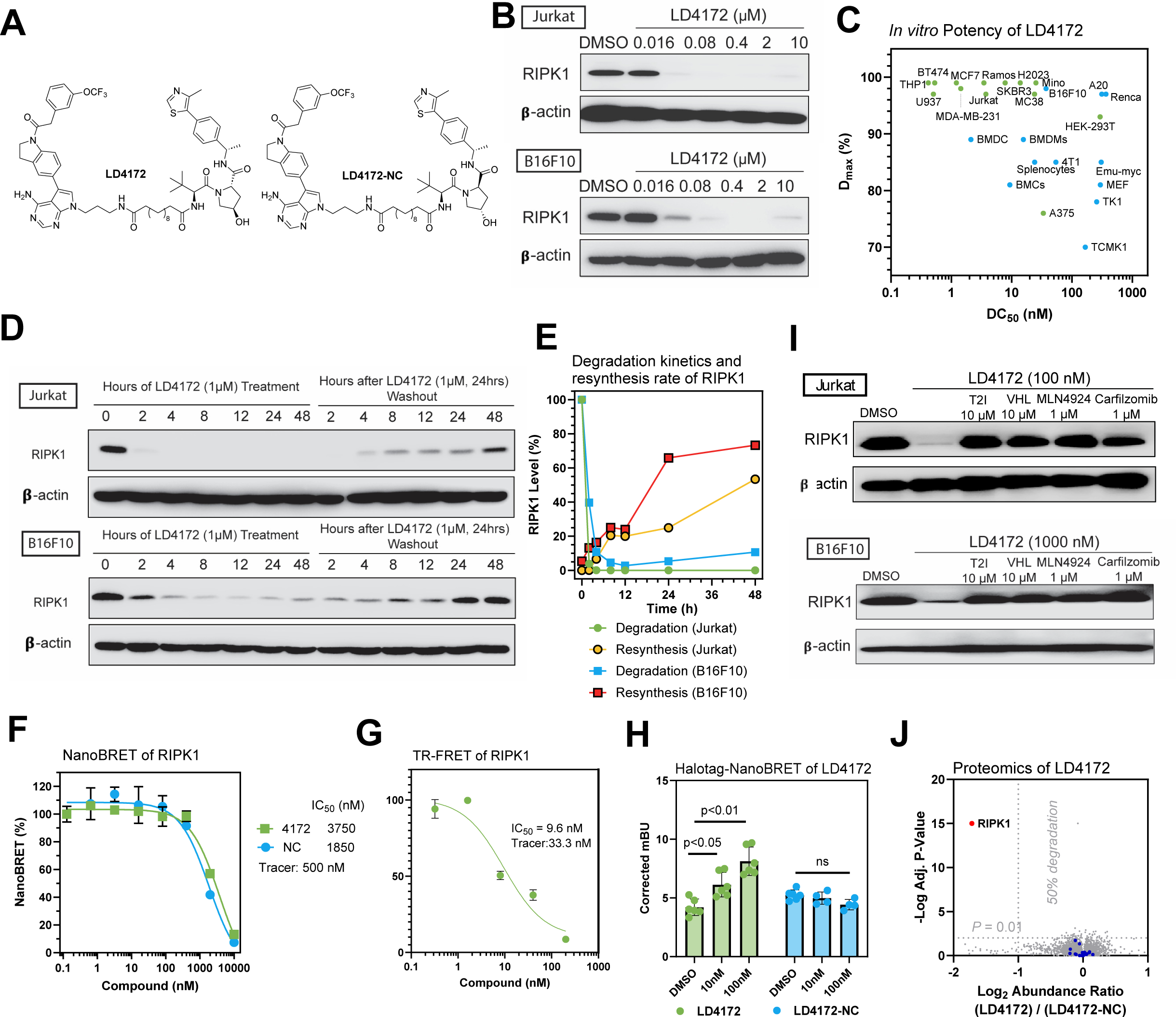
LD4172, a RIPK1 PROTAC, induces potent and highly specific degradation of RIPK1 in a panel of cell lines. **A**, Chemical structures of RIPK1 PROTAC, LD4172, and its negative control, LD4172-NC. LD4172-NC has an identical warhead and linker as LD4172 but with an inactive VHL ligand and is therefore unable to engage VHL to induce ubiquitination. **B**, Quantification of RIPK1 levels in both Jurkat and B16F10 cells treated with LD4172 at indicated concentrations for 24 h, followed by Western blotting. **C**, DC_50_ and D_max_ of LD4172 in various human and mouse cell lines. DC_50_, the drug concentration causing 50% protein degradation; D_max_, the maximum level of protein degradation. **D**, RIPK1 degradation kinetics induced by LD4172 treatment (1 µM) and resynthesis kinetics upon washout of LD4172 in both Jurkat and B16F10 cells. **E**, Quantification of **D**. The degradation half-life of RIPK1 induced by LD4172 (1 µM) is <2 h in Jurkat and B16F10 cells. The resynthesis half-life of RIPK1 is ∼48 h and ∼24 h in Jurkat and B16F10 cells, respectively. **F**, NanoBRET based in-cell RIPK1 target engagement. HEK293 cells were transfected with a plasmid expressing nLuc-RIPK1 fusion protein for 24 h, followed by incubating with a RIPK1 NanoBRET tracer (500 nM) and different concentrations of LD4172 and LD4172-NC (n=3). **G**, Time-resolved fluorescence resonance energy transfer (TR-FRET) based biochemical binding assay for RIPK1. GST-tagged human RIPK1 (1 nM), Tb-labeled anti-GST antibody (0.3 nM), a RIPK1 TR-FRET tracer (350 nM) and different concentrations of LD4172 (n=3) were incubated for 2 hours, followed by TR-FRET measurements with an excitation wavelength at 340 nm and emission wavelengths at 495 and 520 nm. **H**, NanoBRET based in-cell assay for ternary complex formation. HEK293T cells were co-transfected with nLuc-RIPK1 and VHL-HaloTag plasmids for 24 h, followed by treatment with indicated concentrations of LD4172 or LD4172-NC (n=3). **I**, LD4172 induced RIPK1 degradation depends on ternary complex formation, neddylation and proteasome activity. Representative immunoblots of RIPK1 in both Jurkat and B16F10 cells. Cells were treated with RIPK1 inhibitor (T2I), VHL ligand, a neddylation inhibitor (MLN4924) or a proteasome inhibitor (Carfilzomib) at indicated concentrations for 4 hours, followed by LD4172 treatment for 4 hours. **J**, Proteome profiling of LD4172 induced protein degradation. MDA-MB-231 cells were treated with LD4172 (200 nM) or LD4172-NC (200 nM) for 6 h (n=3). In total, ∼10,000 proteins were quantified in the proteomics experiment. RIPK1 (red dot) is the only protein showing >50% degradation with p < 0.01. Blue dots represent kinases that are inhibited by the warhead of LD4172 but not degraded by LD4172.

### LD4172 Induces Potent RIPK1 Degradation *In Vitro*

LD4172 induced potent RIPK1 degradation (concentration to induce 50% protein degradation DC_50_ = 4 to 400 nM) in a panel of human and mouse cancer cell lines (**Fig. 2B-C, S1**). To investigate the kinetics of LD4172-induced RIPK1 degradation and resynthesis rates, Jurkat and B16F10 cells were treated with LD4172 for different time points, followed by washout after 24 h. With 1 µM LD4172 treatment, >90% of RIPK1 was degraded within 2 and 4 h in Jurkat and B16F10 cells, respectively (**Fig. 2D-E**). Upon removal of LD4172, the re-synthesis half-life of RIPK1 was ∼48 h and ∼24 h in Jurkat and B16F10 cells, respectively (**Fig. 2D-E**). Collectively, these data demonstrated that LD4172 is a potent RIPK1 degrader with rapid and sustained effects *in vitro*.

### LD4172 Engages RIPK1 and Forms a Ternary Complex

To elucidate the formation of a binary complex during RIPK1 degradation, we developed a competitive NanoBRET (Nano-Bioluminescence Resonance Energy transfer)-based target engagement (TE) assay to quantify the binding between RIPK1 and LD4172 in cells ^12,13^. First, we developed a RIPK1 tracer by conjugating the LD4172 warhead T2I with a BODIPY-590 fluorescent dye, dubbed T2-590 (refer to Supporting Information for details). The dissociation equilibrium constant (*K*_d_) between the tracer and RIPK1 was determined to be 0.5 µM by titrating the tracer in HEK293T cells expressing a nLuc-RIPK1 fusion protein. Subsequently, with the tracer concentration at its *K*_d_ value, LD4172 competed with the tracer with an IC_50_ value of 3.7 µM (**Fig. 2F**). Based on the Cheng-Prusoff equation, the apparent *K*_i_ between LD4172 and RIPK1 in the cells was 1.9 µM. Using a recombinant human RIPK1 protein, we measured the biochemical *K*_i_ between LD4172 and human RIPK1 to be 4.8 nM (**Fig. 2G**), which is 395 folds smaller than the corresponding *K*_i_ in cells. This is usually expected, considering that the large molecular weight of LD4172 may lead to poor cellular permeability. However, the fact that the DC_50_ values of LD4172 are much smaller than its TE IC_50_ values demonstrates the sub-stoichiometric degradation of RIPK1 induced by LD4172. Additionally, we synthesized an LD4172 negative control (LD4172-NC, also referred to as NC, **Fig 2A**) using a VHL ligand diastereomer that does not bind to VHL. As expected, LD4172-NC showed TE similar to that of LD4172 (**Fig. 2F**).

To test whether LD4172 induces ternary complex formation with RIPK1 and VHL, we co-transfected HEK293 cells with nLuc-RIPK1 and VHL-Halo labeled with BODIPY-590 dye. The addition of LD4172, but not LD4172-NC, induced NanoBRET between RIPK1 and VHL, demonstrating the formation of a ternary complex among {RIPK1-LD4172-VHL} (**Fig. 2H**).

### LD4172 Degrades RIPK1 with High Specificity Through Ubiquitin-Proteasome System (UPS)

The mechanistic action of PROTACs involves bringing the protein of interest (POI) into close proximity to E3 ligase, which ubiquitinates the POI for degradation by the proteasome. To confirm that LD4172 functions through this mechanism, we disrupted ternary complex formation by introducing an excess of RIPK1 or VHL ligands, which led to attenuation of RIPK1 degradation induced by LD4172. Moreover, blocking Cul2 E3 ligase with the neddylation inhibitor MLN4924 or inhibiting the proteasome with carfilzomib reversed the potent degradation of RIPK1 by LD4172 in both Jurkat and B16F10 cells (**Fig. 2I**). These findings indicate that LD4172 induces protein degradation through ternary complex formation and the UPS machinery.

The RIPK1 binder used in LD4172 is a typical type II kinase inhibitor bound to some off-target kinases, including TrkA, Flt1, Flt4, Ret, Met, Mer, Fak, FGFR1, and MLK1 ^10^. To evaluate the specificity of LD4172, we performed mass spectrometry (MS) analysis of the whole cellular proteome. Because Jurkat and B16F10 cells lack expression of all the aforementioned off-target kinases, MDA-MB-231 cells were chosen and treated with either LD4172 (200 nM) or LD4172-NC (200 nM) for 6 h. Among the >10,000 proteins detected, RIPK1 was the only protein degraded by LD4172 (the red dot in **Fig. 2J**), and no degradation of off-target kinases was observed (blue dots in **Fig. 2J**, Supporting data values). This finding is consistent with previous studies showing that PROTACs with promiscuous target protein binders can achieve enhanced selectivity through protein-protein interactions with the E3 ligase involved ^14^.

### LD4172 Sensitizes B16F10 Cells to TNFα-Mediated Apoptosis

In contrast to situations where RIPK1 is kinase-dead, genetic deletion of RIPK1 has been found to trigger apoptosis both *in vitro* and *in vivo*^15^. To investigate apoptosis in the B16F10 mouse melanoma cell model and its correlation with the mechanism of RIPK1 downregulation rather than kinase inhibition, we employed various tool molecules, including LD4172, T2I, TNFα, and the pan-caspase inhibitor Z-VAD-FMK. Results demonstrated that significant cell death (**Fig. 3A-C**), particularly apoptosis, was induced by the combination of TNFα and LD4172, as evidenced by enhanced surface exposure of phosphatidylserine (**Fig. 3A**), along with increased expressions of cleaved caspase3/7 and PARP (**Fig. 3B-C**), which can be reversed with Z-VAD-FMK treatment (**Fig. 3A-C**). In contrast, inhibition of RIPK1 kinase activity by T2I did not trigger TNFα-mediated apoptosis (**Fig. 3A-C**). Additionally, apoptosis induced by LD4172 plus TNFα involves membrane ruptures, as indicated by the enhanced production of ATP in extracellular environments (**Fig. 3D**), loss of nuclear HMGB1 (High Mobility Group Box 1, **Fig. 3D-E**) and downregulated calreticulin (**Fig. 3D**).

**Figure 3.**
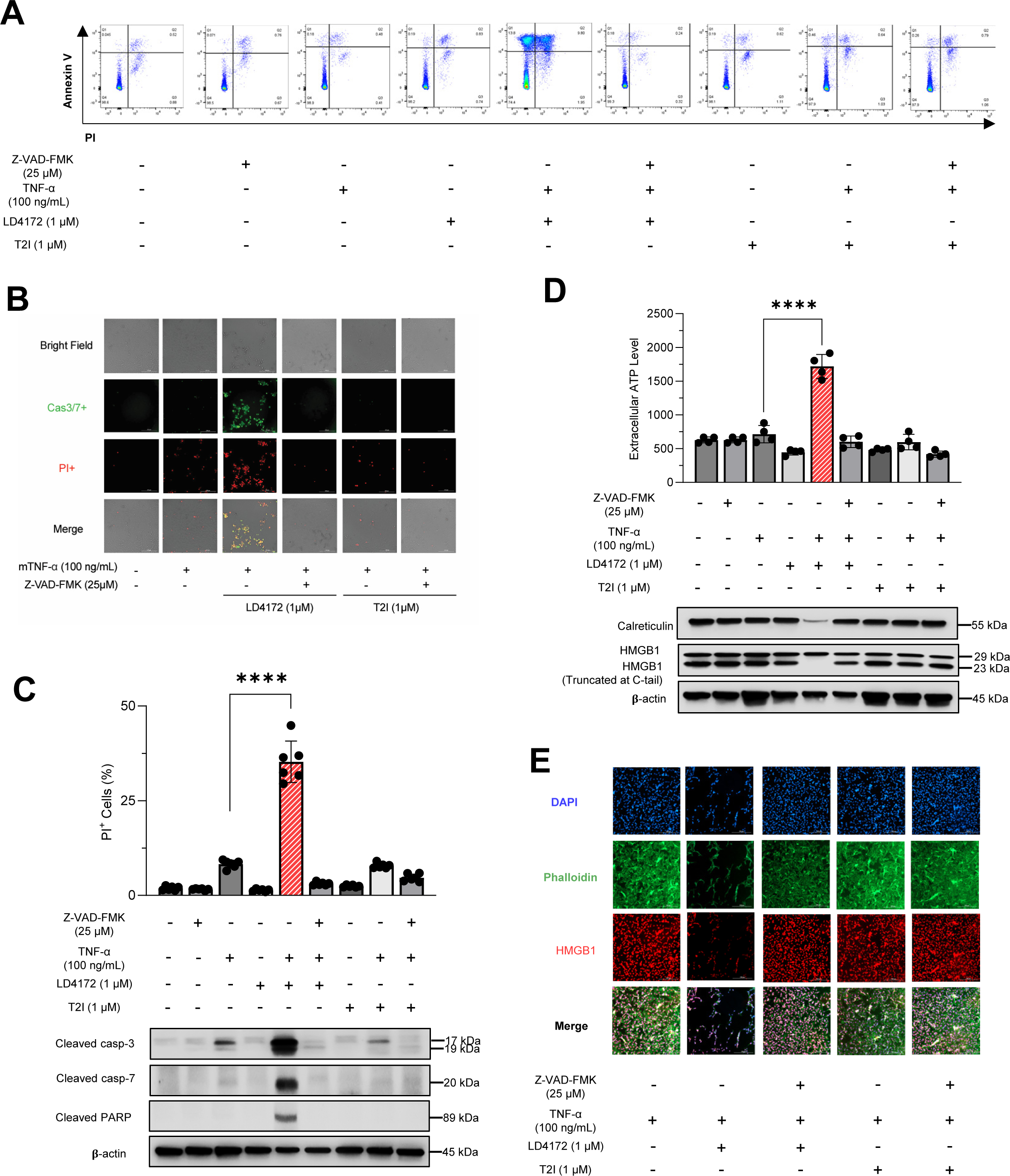
LD4172 sensitizes B16F10 cells to TNFα-mediated apoptosis. **A**, Representative flow cytometry dot plots of apoptosis. In all four plots, viable cells are seen in the left lower quadrant (FITC-/PI-), early apoptotic cells in the right lower quadrant (FITC+/PI-), and late apoptotic cells in the right upper quadrant (FITC+/PI+). **B**, Representative images of PI (red) and caspase 3/7 (Cas3/7+, green) staining of B16F10 cells. Cell death assessed by PI uptake using Cytation 5 imager. **C**, Western blots for the expression of cleaved caspase3, cleaved caspase7, and cleaved PARP in B16F10 cells. **D**, Quantification of extracellular ATP level. Western blots for the expression of HMGB1 and calreticulin in B16F10 cells. **E**, Immunofluorescence staining of HMGB1 in B16F10 cells. B16F10 cells were treated in the presence or absence of TNFα (100 ng/mL), Z-VAD-FMK (25 μM), LD4172 (1 μM), and/or T2I (1 μM) as indicated for 72 hours. Data are representative of three independent experiments. Error bars represent SD.

### LD4172 Exhibits Acceptable Pharmacokinetic Properties and Tissue-selective RIPK1 Degradation

LD4172 has half-lives of 21.1 and 9.7 minutes in human and mouse liver S9 fractions, respectively, corresponding to intrinsic clearance (CL_int_) of 32.8 and 71.6 µL·min^-1^·mg^-1^ protein. In human primary hepatocytes, the half-life of LD4172 is 56.3 minutes, which corresponds to a predicted CL_int_ of 15.6 mL·min^-1^·kg^-1^ in human. It should be noted that the predicted intrinsic clearance in primary hepatocytes was >2,000 times slower than that in liver S9 fractions, possibly due to the low membrane permeability of LD4172, which protects it from being metabolized (**Table 1**).

**Table 1:**
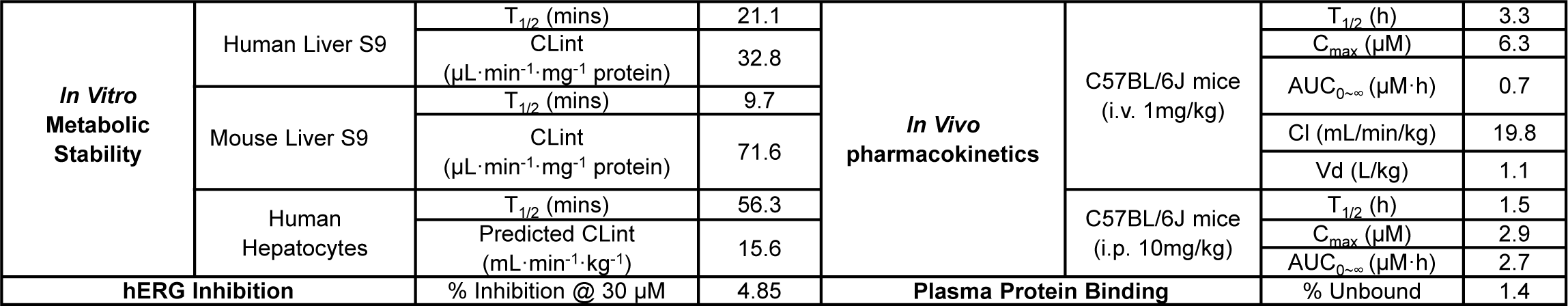
Absorption, distribution, metabolism, excretion, and toxicity (ADMET) summary of LD4172.

Next, we evaluated the pharmacokinetics (PK) of LD4172 in C57BL/6J (B6) mice (**Fig. 4A**). With 1 mg/kg intravenous (i.v.) administration in C57BL/6J mice, LD4172 showed a half-life (t_1/2_) of 3.3 ± 2.1 h, a maximum plasma concentration (C_max_) of 6.3 ± 0.8 µM, and an area under the concentration-time curve (AUC) of 0.7 ± 0.07 µM·h. The volume of distribution (V_d_) of LD4172 was 1100 ± 200 mL·kg^-1^, which is much greater than the mouse plasma volume (77-80 mL·kg^-1^), suggesting that LD4172 has strong affinities to tissues. The clearance of LD4172 is 19.8 ± 1.8 mL·min^-1^·kg^-1^.

**Figure 4.**
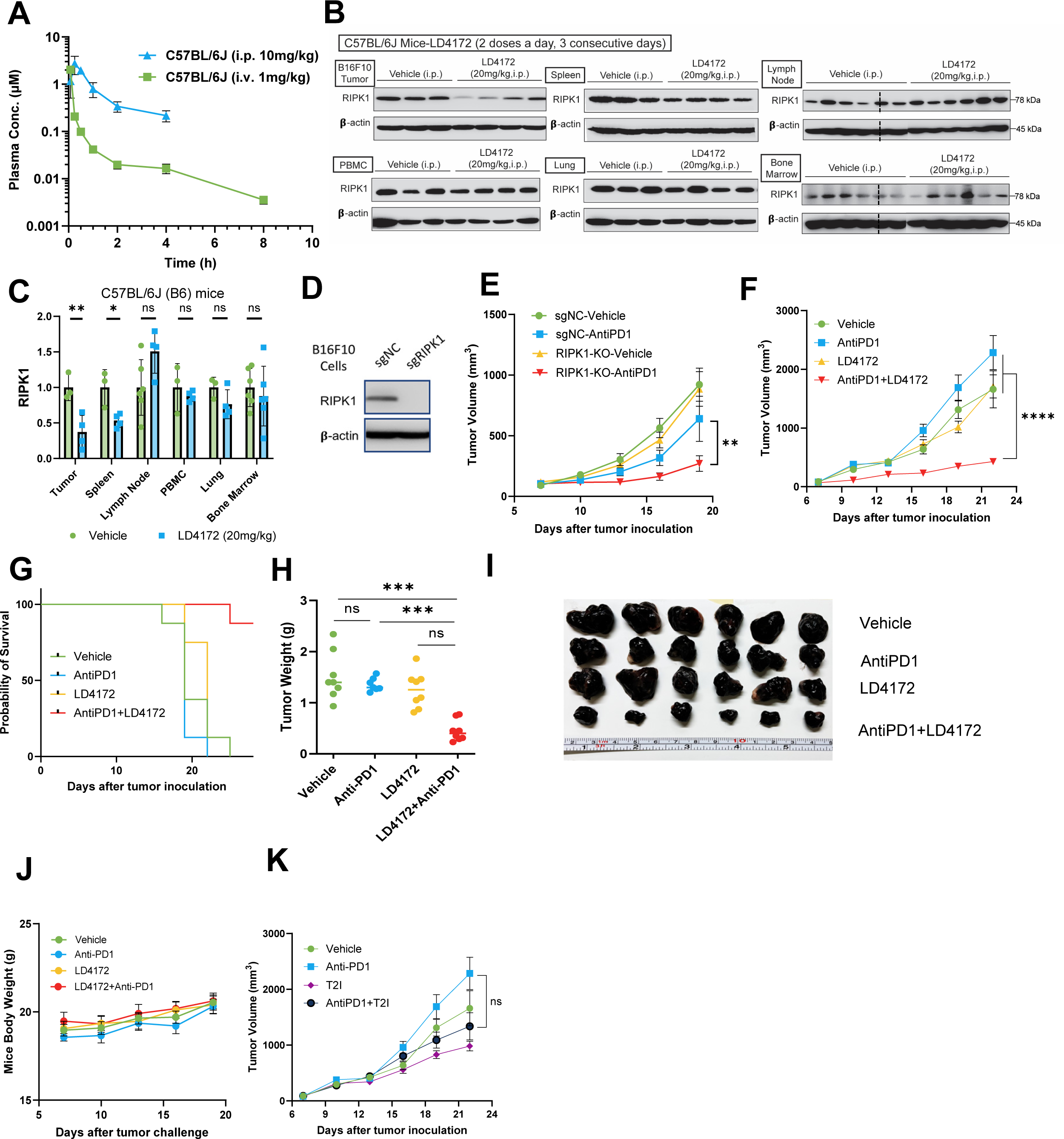
LD4172 synergizes with Anti-PD1 to inhibit tumor growth. **A**, Plasma concentrations of LD4172 in C57BL/6J mice administered 1 mg/kg intravenously (i.v.) or 10 mg/kg intraperitoneally (i.p.) (n=4). **B**, Representative immunoblots of RIPK1 in different tissues of C57BL/6J mice treated with LD4172 (n=3-4). **C,** Densitometric analysis of RIPK1 protein levels in different tissues (n=3 or 4). **D**, Representative immunoblots of RIPK1 levels of B16F10 cells transduced with gRNA specific for RIPK1 or non-targetable control (sgNC). **E,** B16F10-RIPK1-KO tumors sensitized to anti-PD1 treatment: 3×10^5^ B16F10 cells transduced with gRNA specific for RIPK1 or non-targetable control (sgNC) were inoculated into C57BL/6J mice. After seven days, mice with measurable tumors (∼100 mm^3^) were randomly treated with or without anti-PD1 *in vivo* (100 µg per dose, i.p., every three days, n=8). **F**, Tumor growth curve of mice with B16F10 tumors treated with LD4172 and/or anti-PD1 (n=8). C57B6/J mice were subcutaneously inoculated with 3×10^5^ B16F10 tumor cells. After seven days (tumor size ∼ 100 mm^3^), mice were treated every three days with anti-PD1 (100 µg per dose, i.p.), daily with LD4172 (20 mg/kg, i.p.), a combination of LD4172 and anti-PD1 (same dose as their individual doses), or their corresponding vehicle control. **G**, Kaplan-Meier survival curve for all experimental groups. **H**, Final tumor weight (g) from **F** after 22d of treatment (n=8). **I**, Representative images of B16F10 tumors collected at the end of treatment. **J**, Mouse body weight (n=8). **K,** Tumor growth curve of mice with B16F10 tumors treated with T2I and/or anti-PD1 (n=8). The experimental conditions and treatment regimens were the same as **F** except using the RIPK1 kinase inhibitor T2I (20 mg/kg, i.p.) to replace LD4172. Representative data from at least three independent experiments are shown. Data are expressed as the mean ± SEM. * p<0.05; ** p<0.01; *** p<0.001, **** p<0.0001, respectively. ns, indicates no statistical significance.

Intraperitoneal (i.p.) administration of LD4172 (10 mg/kg) to C57BL/6J mice led to C_max_, t_1/2_, and AUC as 2.9 µM, 1.5 h, and 2.7 µM·h, respectively (**Fig. 4A**, **Table 1**). Considering an AUC of 0.7 µM·h for i.v. administration (1 mg/kg), i.p. administration of LD4172 achieved 39% bioavailability.

To investigate the pharmacodynamics of LD4172 *in vivo*, we administered LD4172 via the i.p. route and observed a 60% reduction in RIPK1 levels in tumors (20 mg/kg, b.i.d., i.p.) (**Fig. 4B-C**). In contrast, less than 50% RIPK1 degradation was observed in the spleen, and no significant RIPK1 degradation was observed in other organs, including the lymph nodes, PBMCs, lungs, and bone marrow (**Fig. 4B-C**).

The hERG channel inhibition assay is a commonly used safety assay to identify compounds that exhibit cardiotoxicity related to hERG inhibition *in vivo*. LD4172 exhibited no obvious inhibition of hERG, even at 30 µM (**Table 1**), indicating that LD4172 has a good safety margin for hERG inhibition.

### LD4172 Sensitizes Tumors to Anti-PD1 Therapy

Utilizing CRISPR-Cas9 technology, we generated RIPK1-knockout (KO) B16F10 cells and implanted them into mice to examine their response to anti-PD1 treatment (**Fig. 4D-E**). Align with previous reports^5–7^, our findings demonstrated that tumors lacking RIPK1 exhibit heightened sensitivity to anti-PD1 treatment (**Fig. 4E**). Subsequently, we explored whether pharmacological degradation of RIPK1 could replicate the effects observed in RIPK1-null B16F10 tumors. Consistent with the genetic study, mice treated with anti-PD1 or LD4172 alone showed tumor progression similar to that of the untreated mice. However, LD4172 sensitized B16F10 tumors to anti-PD1 therapy (**Fig. 4F-I**), with long-term administration of LD4172 showing no impact on mouse body weight (**Fig. 4J**). To test whether inhibition of RIPK1 kinase activity also enhances tumor responses to ICB therapy, we treated B16F10 xenograft tumors with the RIPK1 kinase inhibitor T2I, alone or in combination with anti-PD1. Unlike the RIPK1 degrader LD4172, the RIPK1 kinase inhibitor T2I failed to sensitize B16F10 tumors to anti-PD1 treatment (**Fig. 4K**).

We also tested a syngeneic MC38 colon cancer model, which exhibited a limited response to anti-PD1 treatment. Consistent with the B16F10 tumor model, LD4172 substantially sensitized MC38 tumors to anti-PD1 therapy (**Fig. S2A-B**).

### LD4172 Triggers Immunogenic Cell Death in B16F10 Tumor

To understand the observed synergistic effects of LD4172 and anti-PD1, we administered vehicle, LD4172, anti-PD1, or a combination of LD4172 and anti-PD1 in C57BL/6J mice with B16F10 tumors for a short duration. A five-day treatment with LD4172 was sufficient to induce substantial degradation of RIPK1 in the tumor (**Fig. 5A, 1^st^ column**). Consistent with the *in vitro* findings, LD4172 also triggered significant cell death in the tumor (**Fig. 5A, 2^nd^ column**). Importantly, a notable increase in cleaved caspase 3/7 levels was observed in the LD4172-treated tumors, indicating the occurrence of apoptosis (**Fig. 5A, 3^rd^ and 4^th^ columns**). While apoptotic cell death was traditionally considered non-immunogenic, accumulating experimental data have revealed its potential to drive immune cell infiltration and anti-cancer immunity ^16–19^. Supporting the activation of immunogenic apoptosis, we observed a significant increase in plasma HMGB1 levels (**Fig. 5B**) and enhanced exposure of calreticulin on the surface of B16F10 tumor cells (**Fig. 5A, 5^th^ column**).

**Figure 5.**
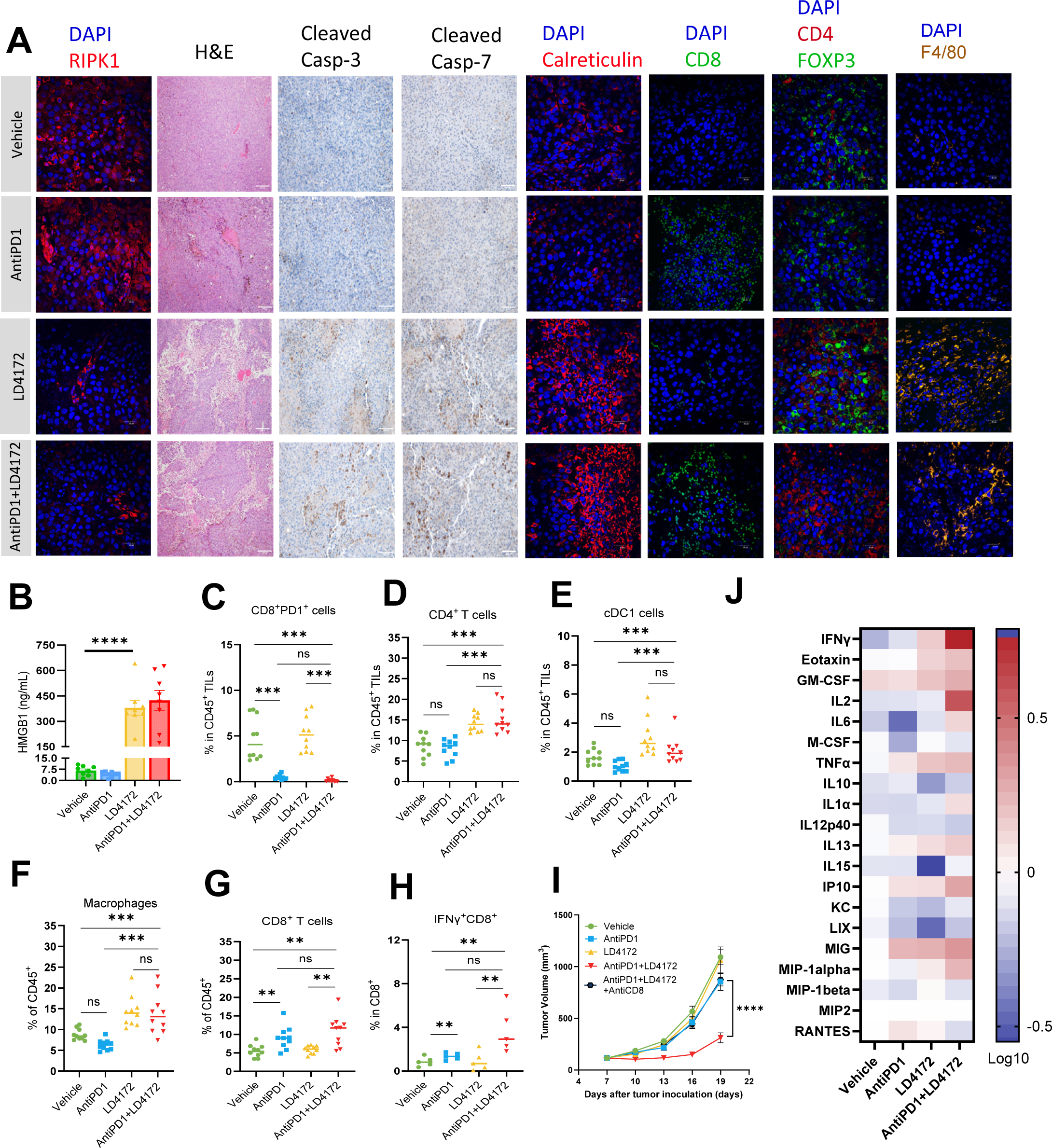
LD4172 alters the tumor immune microenvironment. **A.** Representative immunofluorescent images depict RIPK1, Calreticulin, CD8, CD4, Foxp3, or F4/80, as well as hematoxylin and eosin (H&E) images, alongside representative images illustrating cleaved caspase3/7 levels in B16F10 tumors under the indicated treatment for five days. n = 4-6 independent fields per group. Scale bars are shown as follows: 20 μm, objective 60X; 200 μm, objective 60X. **B**. The mouse plasma HMGB1 level from different treatment groups (n=8). **C-G**. Flow cytometry quantification of PD1+CD8+ T cells (**C**), CD4+ T cells (**D**), cDC cells (**E**), macrophages (**F**), and CD8+ T cells (**G**) in B16F10 tumors following 5 days of indicated treatment (n = 10/group). **H**. Flow cytometric quantification of IFNγ+ CD8+ T cells in B16F10 tumors treated with the indicated treatments for 5 days and stimulated with PMA/ionomycin *in vitro* for 6 h (n = 10/group). **I**. Tumor volume (mm^3^) of vehicle-, anti-PD1-, LD4172-, anti-PD1+LD4172-, and anti-PD1+LD4172+anti-CD8–treated B16F10 tumors (n=8). **J**. Heat map showing log 10-fold changes in the concentration of mouse plasma cytokines normalized by the mean value of control mice (n=8). For all experiments: LD4172: 20 mg/kg; anti-PD1 antibody: 100 μg/mice; Combo: LD4172 plus anti-PD1. Representative data from at least three independent experiments are shown. Data are expressed as the mean ± SEM. * p<0.05; ** p<0.01; *** p<0.001; **** p<0.0001, respectively. “ns” indicates no statistical significance.

### LD4172 Enhances Anti-tumor Immunity

To elucidate how the combination of LD4172 plus anti-PD1 promotes anti-tumor immunity, multiparameter flow cytometry was employed to evaluate tumor-infiltrating lymphocytes (TILs) within the tumor microenvironment (TME) of mice receiving different treatments (**Fig. S3**). Initially, we confirmed the successful blockade of PD1 on T cells (CD8+PD1+) with an anti-PD1 antibody (**Fig. 5C**). LD4172-induced ICD led to a notable expansion of CD4+ T cells (**Fig. 5A, 7^th^ column, and 5D**), conventional dendritic cells (cDC1, CD45+CD11C+IAIE+XCR1+, **Fig. 5E**), and macrophages (CD45+CD11b+F4/80+, **Fig. 5A, 8^th^ column, and 5F**) within the TME, all of which contribute to antigen presentation and cytotoxic T cell priming and activation. In addition, combined therapy with LD4172 and anti-PD1 not only induced extensive TIL infiltration (**Fig. 5D-H**) but also significantly enhanced anti-PD1 positivity in immunologically cold B16F10 tumors, as demonstrated by increased infiltration of cytotoxic CD8+ T cells (CD8+IFN-γ+, **Fig. 5A, 6^th^ column, and 5G-H**) and decreased infiltration of FOXP3+ T regulatory cells (**Fig. 5A, 7^th^ column**) within the TME. Additionally, to confirm the contribution of CD8+ T cells to the antitumor effect, we conducted a CD8+ T cell depletion experiment, revealing that the synergy between anti-PD1 and LD4172 was nullified in the absence of CD8+ T cells (**Fig. 5I**). Results from the cytokine array profiling of plasma further supported synergistic effects of combined treatment, showing a significant enhancement in the production of immune cell proliferation cytokines, including IFN-γ and IL2 (**Fig. 5J**).

## Discussion

RIPK1 is a critical regulator involved in cellular processes and proinflammatory signaling and exerts its effects through both kinase-dependent and kinase-independent mechanisms. In particular, its ubiquitin scaffolding function through K376 has been implicated in conferring intrinsic and extrinsic resistance to immune checkpoint blockade and is a potential target for cancer immunotherapy ^5,6^. However, the development of inhibitors that specifically target the intermediate scaffolding domain is challenging because of the absence of a well-defined binding pocket. In this study, we used PROTAC technology to address this limitation and successfully developed a first-in-class RIPK1 degrader, LD4172, with potent and specific RIPK1 degradation both *in vitro* and *in vivo*. Notably, LD4172 also exhibited therapeutic efficacy *in vivo*, leading to RIPK1 degradation in tumors and demonstrating a synergistic effect in inhibiting tumor growth when combined with anti-PD1 treatment. These findings highlight the potential of developing RIPK1 degraders as a promising therapeutic strategy to enhance the antitumor immunity of anti-PD1 blockades.

The suboptimal pharmacokinetic properties of LD4172, particularly its high *in vivo* clearance and low unbound plasma drug concentration, present challenges for achieving optimal RIPK1 degradation *in vivo* following intraperitoneal administration. Interestingly, intratumoral administration of LD4172 improved RIPK1 degradation in the tumors (**Fig. S4**), suggesting that the incomplete degradation of RIPK1 in tumors via i.p. injections may be attributed to poor penetration and/or accumulation of LD4172 in tumor tissues. Additionally, LD4172 had poor permeability in cells based on the divergence observed between biochemical and cellular assays for target engagement (**Fig. 2G-H**). To enhance the pharmacodynamic properties of LD4172, optimization of medicinal chemistry is necessary to further improve its physicochemical and pharmacokinetic characteristics. Previous studies have reported successful optimization strategies, including optimizing linker structures ^20^, E3 ligands, and kinase warheads ^21^, introducing intramolecular hydrogen bonds ^22^, and converting the drug into a prodrug form ^23^. Our future work will implement these optimization approaches to enhance membrane permeability and the overall pharmacokinetic profile of LD4172, thereby improving its potency for RIPK1 degradation *in vivo*.

As a first-in-class therapeutic modality, the on-target toxicity profiles of RIPK1 degraders remain unclear. Although mice are a convenient model system for exploring the functions of cellular signaling pathways, human genetics provides the best models of human diseases and guides the selection of new targets for drug discovery ^24^. Unlike *Ripk1* knockout mice, which die at 1-3 days of age due to their widespread roles in multiple tissues and organs ^25^, homozygous loss-of-function *RIPK1* mutations are well tolerated in humans ^26^. Patients with complete loss of RIPK1 protein only showed symptoms confined to the immune system, with primary immunodeficiency and/or intestinal inflammation ^26^. Additionally, the phenotypes of genetic knockout may be different from those of chemical-induced protein degradation, which is acute and transient and can be tissue-specific ^27^. Although the safety profiles of RIPK1 degraders remain to be tested in future clinical studies, human genetic data suggest that pharmacological RIPK1 degradation is potentially safe and tolerable, especially with transient intervention in well-controlled clinical settings. Furthermore, we found that decreasing RIPK1 dosing frequency by 50% resulted in a similar tumor inhibition effect when combined with anti-PD1 (**Fig. S5**), suggesting that the safety profile of RIPK1 degraders can be further improved by optimizing the dosing regimen.

Importantly, we primarily observed RIPK1 degradation in tumors to a lesser extent in the spleen and no degradation in other organs (**Fig. 4B-C**). In contrast to small-molecule inhibitors, protein degraders can achieve tissue-specific target degradation by leveraging tissue-specific E3 ligases. One notable example is a BCL-X_L_ degrader developed by the groups of Zhou and Zheng, which spares platelets due to the low expression of VHL in platelets ^28^. However, according to proteinatlas.org, VHL expression is ubiquitous in major organs, which does not explain the tumor-selective RIPK1 degradation induced by LD4172. Albumin accounts for ∼60% of the total plasma protein and preferentially accumulates in tumors due to the high demand for amino acids and energy ^29,30^. Considering that 98.6% of LD4172 is bound to plasma proteins (**Table 1**), it is possible that LD4172 piggybacks albumin accumulation in tumors to achieve tumor-selective RIPK1 degradation, further alleviating potential toxicity concerns associated with RIPK1 degradation in normal tissues.

In summary, we developed a first-in-class RIPK1 degrader with high degradation specificity in cells and tumor selectivity *in vivo*. Our work not only provides a high-quality chemical probe to explore the effects of RIPK1 degradation in biology, but also the first proof-of-concept study demonstrating that pharmacological degradation of RIPK1 synergizes with anti-PD1 to overcome resistance to ICBs by enhancing the infiltration of effector immune cells and promoting the secretion of immunostimulatory cytokines. Considering the predicted safety profile of RIPK1 degradation based on human genetics, we envision that further optimized RIPK1 degraders have the potential to improve cancer immunotherapy.

### Methods Cell Lines

Human and mouse hematopoietic cell lines, namely Jurkat, Ramos, THP1, U937, TK1, and A20, and mouse melanoma B16F10 cell lines, were procured from ATCC. MC38 colon carcinoma and H2023 lung carcinoma cells were provided by Dr. Weiyi Peng. A375 melanoma cells were acquired from the Cell Core at the MD Anderson Cancer Center. Human breast cancer cells MDA-MB-231 and BT474 were obtained from Baylor College of Medicine Cell Core, whereas mouse breast carcinoma 4T1 cells were a gift from Dr. Xiang Zhang.

The suspension cell lines were cultured in RPMI-1640 medium (MT10040CV, Thermo Fisher Scientific), whereas adherent cell lines were cultured in Dulbecco’s modified DMEM medium (MT10013CV, Thermo Fisher Scientific). Both media were supplemented with 10% fetal bovine serum (SH30071.03, GE Healthcare), and 1% penicillin-streptomycin (15140163, Thermo Fisher Scientific). All cell lines were maintained in a humidified incubator at 37 °C with 5% CO_2_.

### Induction of Bone Marrow-Derived Macrophages and Dendritic Cells

To isolate bone marrow cells, femur and tibia bones were dissected from 6-to 8-week-old C57BL/6J mice. The bone marrow was flushed out into cold PBS containing 2% heat-inactivated fetal bovine serum using a 27G needle and 1 mL syringe. The cells were dissociated by passing the bone marrow through a 70µm cell strainer and then incubated in RBC lysis buffer (1X, 420301, Biolegend) on ice for 10 minutes. After centrifugation (500×g, 5 minutes, 4°C), the supernatant was discarded, and the cells were resuspended in 1 mL of BMDM/BMDC growth medium. A suspension of 1×10^6^ cells was prepared in 20 mL of cell culture medium and plated into a 10 cm petri dish. Subsequently, BMDM were induced with G-CSF (25 ng/mL, 250-05, Peprotech), and BMDCs were induced with GM-CSF (20 ng/mL, 315-03, Peprotech) and IL4 (10 ng/mL, 214-14, Peprotech).

### Western Blotting

Cells were seeded into six-well plates at a density of 5×10^5^ cells/mL in 2 mL of complete culture medium. Following an overnight adaptation period, cells were treated with serially diluted LD4172 compounds for 24 h. After treatment, whole-cell lysates were prepared using a lysis buffer (1×RIPA supplemented with protease and phosphatase inhibitor cocktail). Protein concentrations in the lysates were measured using the BCA protein assay. Subsequently, equal amounts of protein (20 µg) from each sample were loaded onto a sodium dodecyl sulfate-polyacrylamide gel and separated by electrophoresis (Bio-Rad) at 120 V for 1.5 hours. The separated proteins were then transferred to a polyvinylidene fluoride (PVDF) membrane using a Transblot Turbo system (Bio-Rad). After blocking for 1 h at room temperature in 1% BSA-TBST, the membranes were incubated overnight at 4°C with specific primary antibodies (diluted at 1:1000 in TBST) targeting the proteins of interest, including anti-RIPK1 (3493, Cell Signaling Technology (CST)), anti-cleaved caspase 3 (9661, CST), anti-cleaved caspase7 (8438, CST), anti-cleaved PARP (5625,CST), anti-HMGB1, anti-calreticulin, and anti-β-actin (4970, CST). The membranes were then incubated with horseradish peroxidase-conjugated secondary antibodies (1:1000, 7074, CST) for 1 h at room temperature. Immunoblots were imaged using ECL Prime chemiluminescent western blot detection reagent (Kindle Biosciences, Cat. No. R1100) and visualized using an Imager (Kindle Biosciences, Cat. No. D1001). All western blots were processed and quantified using ImageJ software, and protein levels were normalized to β-actin loading controls.

### Apoptosis detection using FITC-conjugated Annexin V/PI

Apoptosis quantification was conducted utilizing a FITC-conjugated Annexin V/PI assay kit (BD, 556547) and analyzed through flow cytometry. Briefly, 2×10^5^ of B16F10 cells were seeded onto six-well plates and treated as specified for 72 hours at 37°C. Treated and untreated cells were harvested, washed with PBS, and resuspended in 500 µl of binding buffer. Subsequently, cells were stained with PI (50 µg/ml) and FITC-conjugated Annexin V (10 mg/ml) for 15 minutes at room temperature in the dark. Analysis was performed using a BD-LSR II Flow cytometer (BD Biosciences), and flow cytometry data were processed using the FlowJo software.

### Extracellular ATP Assay

Extracellular ATP (eATP) is a potent damage-associated molecular pattern (DAMP) molecule known to exert profound effects on innate and adaptive immune responses. To detect ATP secretion after treating B16F10 cells with specified treatments, the RealTime-Glo™ Extracellular ATP Assay (GA5010, Promega) was conducted following the manufacturer’s protocol. In brief, 1×10^4^ B16F10 cells were plated into each well of an opaque 96-well plate, after 72 hours of treatment, 1X assay reagent was dispensed, and luminescence was recorded at regular intervals.

### TR-FRET Biochemical Binding Assay

A time-resolved fluorescence resonance energy transfer (TR-FRET) assay was performed to evaluate the binding of the indicated compounds and RIPK1 by competition with a BODIPY-FL labeled RIPK1 tracer (Supplementary Information, **T2I-488**). The assay was performed in 20 μL assay buffer (50 mM Tris, pH7.5, 0.1% Triton X-100, 0.01% BSA, and 1mM DTT) with 0.3 nM Tb-anti-GST (61GSTTLF, Cisbio), 2 nM GST-RIPK1 (R07-11G-10, SignalChem), 150 nM RIPK1 tracer, and serially diluted compounds (10,000 to 0.64 nM, 5-fold dilutions) in opaque 384-well plates. Unless specified otherwise, all assays were performed in triplicate. The assay mixtures were incubated at room temperature in the dark for 120 min, and the signals were collected using a BioTek Synergy H1 microplate reader to measure the fluorescence emission ratio (I520 nm/I490 nm) of each well using a 340-nm excitation filter, a 100-μs delay, and a 200-μs integration time. Raw data from the plate reader were used directly for the analysis. The curve-fitting software GraphPad Prism 9 was used to generate graphs and curves and determine IC_50_ values.

### NanoBRET Live-cell Ternary Complex Assay

Human RIPK1 cDNA insert was cloned into pLenti6.2-ccdB-nLuc plasmid (87075, Addgene, a kind gift from Taipale Lab) using gateway cloning kit (11791020, Thermo) and standard protocol to obtain pLenti6.2-RIPK1-nLuc fusion vector. The day before transfection, 1 million HEK293T cells were plated in a 60 mm dish and allowed to grow overnight in DMEM/10% FBS. The next day, the cells were co-transfected overnight at 37 deg with 1ng/ml pLenti6.2-RIPK1-nLuc fusion vector, 100ng/ml HaloTag®-VHL Fusion Vector (N273A, Promega) along with 1ug/ml carrier DNA vector (E4881, Promega) using the calcium phosphate method. After 18h, transfected cells were trypsinized and resuspended in Opti-MEM (11058-021, Gibco) supplied with 4% FBS and 100nM HaloTag® NanoBRET™ 618 Ligand (G9801, Promega) to a cell-density of 0.2M/ml (for background subtraction group, 618 ligand was omitted). Plate 100ul cells into each 96-well (136101, Thermo). The plate was further incubated at 37 °C overnight to allow HaloTag-VHL to be labeled with 618 ligand. Next day, cells were further treated with 10uM MG132 for 0.5h followed by 1μM PROTAC or DMSO for 4h. Immediately before reading the plate, prepare 4x concentrated NanoGlo nLuc substrate (N157, Promega) was prepared by diluting the stock into Opti-MEM 1,000-fold, and the NanoGlo substrate was then added into each well to bring the final concentration to 1x. Donor emission at 450 nM and acceptor emission at 610 nM were measured on a BioTek Synergy H1 plate reader equipped with filter cube set 450/80 and 610 LP. The corrected mBU was calculated as follows:

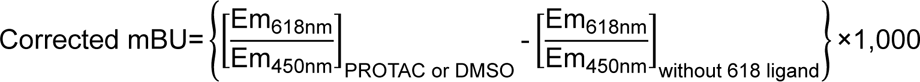

### Proteomics Study

One million MDA-MB-231 cells were seeded in 6-well plates. The following day, cells were treated in triplicate with LD4172 (200 nM) or LD4172-NC (200 nM) for 6 h. The cells were washed thrice with ice-cold PBS. The cell pellets were lysed, reduced, alkylated, and digested using EasyPep™ MS Sample Prep Kits (A45733,ThermoFisher) according to the manufacturer’s instructions. The same amount of peptide from each condition was labeled with a tandem mass tag (TMT) reagent (90113, ThermoFisher). The 10-plex TMT reagent was incubated with each peptide sample at a ratio of 1:8 (peptide:TMT label). The 10-plex labeling reactions were performed for 1 h at room temperature. The labeled peptide samples were quenched by adding 50 µL of 5% hydroxylamine and 20% formic acid solution for 5 min and then mixed. The mixed samples were desalted and fractionated offline into 24 fractions on a 250×4.6 mm Zorbax 300 Extend-C18 column (Agilent) using an Agilent 1260 Infinity HPLC system.

The 24 fractions were dried *in vacuo* and resuspended in 5% acetonitrile in water (0.1% FA). Each sample was first separated by nano LC through a 5-40% ACN gradient within 75 min and ionized by electrospray (2.4 kV), followed by MS/MS analysis using higher-energy collisional dissociation (HCD) at a fixed 38.0 collision energy on an Orbitrap Fusion Lumos mass spectrometer (Thermo Fisher Scientific) in data-dependent mode with a 3 sec cycle-time. MS1 data were acquired using the FTMS analyzer in profile mode at a resolution of 120,000 over a range of 400–1,600 m/z. Following HCD activation and quadrupole isolation with a window of 0.7 m/z, MS/MS data were acquired using an orbitrap at a resolution of 50,000 in centroid mode and normal mass range.

Proteome Discoverer 2.4 (Thermo Fisher Scientific) was used. RAW file processing and controlling peptide- and protein-level false discovery rates, assembling proteins from peptides, and protein quantification from peptides. Searches were performed using full tryptic digestion against the SwissProt human database with up to two miscleavage sites. Oxidation (+15.9949 Da) of methionine and Deamidation on N and Q (0.984 Da) were set as variable modifications, while carbamidomethylation (+57.0214 Da) of cysteine residues and TMT 10-plex labeling of peptide N-termini and lysine residues were set as fixed modifications (+229.163 Da). Data were searched with mass tolerances of ±10 ppm and 0.02 Da on the precursor and fragment ions (HCD), respectively. The results were filtered to include peptide spectrum matches (PSMs) with a high peptide confidence. PSMs with precursor isolation interference values > 50% and average TMT-reporter ion signal-to-noise values (S/N) < 10 were excluded from quantitation. Isotopic impurity correction and TMT channel normalization, based on the total peptide amount, were applied. Protein quantification uses both unique and random peptides. For statistical analysis and adjusted p-value calculation, an integrated analysis of variance (ANOVA) hypothesis test on individual proteins was used. TMT ratios with adjusted p-values below 0.01 were considered significant. The mass spectrometry raw data files for quantitative multiplexed proteomics have been deposited in the MassIVE dataset under accession number *MSV000092377*.

### Molecular Docking

Molecular docking studies were carried out using Schrödinger software. Schrödinger adopted the Glide algorithm to dock flexible ligands into the protein-binding site. The crystal structure of the RIPK1 kinase domain in complex with the isoquinolin-1amine analog (PDB: 4NEU) was used as the receptor structure in molecular docking studies.

### Generation of CRISPR Edited Tumor Cell Lines

RIPK1 genetic deletion was accomplished using the Neon transfection system (ThermoFisher). Five nM of sgRNA targeting mouse RIPK1 (sgRIPK1 #1: GGGTCTTTAGCACGTGCATC, sgRIPK1 #2: CAGTCGAGTGGTGAAGCTAC) or non-targeting negative control sgRNA (sgNC #1: GAAGATGGGCGGGAGTCTTC) was mixed with 2 nM Cas9 enzyme (IDT) at room temperature for 15 min to generate the RNP complex, which was then electroporated into 4 ×10^5^ B16F10 cells. The medium was replaced 24 h after electroporation. Single-cell clones were screened for protein expression by western blotting. Confirmed gene-deleted clones were pooled and cultured for two weeks in vitro before being implanted in vivo.

### Animal studies

All animal experiments were conducted according to the protocol approved by the Institutional Animal Care and Use Committee of the Baylor College of Medicine. All tumor studies were performed in female mice. Female 6-week-old C57BL/6J mice were ordered from Jackson Labs, and experiments were carried out in age-matched animals. Mice were housed in the TMF Mouse Facility at the Baylor College of Medicine under SPF/climate-controlled conditions with 12-hour day or night cycles. They were continuously supplied with fresh chow and water from an autowater system.

### Animal Treatment and Tumor Challenges

To establish a syngeneic mouse model, 3× 10^5^ B16F10 cells resuspended in PBS were mixed 1:1 by volume with Matrigel (354262, Corning) and subcutaneously injected into the right flank of seven-week-old wild-type female C57BL/6J mice on day 0. Treatments were started on day 7 when the tumor volume reached approximately 100 mm^3^. Antibodies were administered in 100μL volumes and injected intraperitoneally every three days for the following: anti-PD1 (100 μg, BE0146, BioXcell) and anti-CD8 (500 µg, BE0004, BioXcell). The RIPK1 degrader LD4172 and RIPK1 kinase inhibitor T2I were delivered in 200 μL of vehicle solution (30% PEG400, 5% Tween-80, 5% DMSO) once daily (20 mg/kg, i.p. injection). T-cell blocking or depletion was confirmed by flow cytometry in tumor-bearing mice. Tumors were measured every three days beginning on day 7 after the challenge until death. Death was defined as the point at which a progressively growing tumor reached 1.5 cm in the longest dimension, or severe tumor necrosis was observed. Measurements were performed manually by collecting the longest dimension (length) and the longest perpendicular dimension (width). Tumor volume was estimated using the following formula: (L×W^2^) / 2. CO_2_ inhalation was used to euthanize mice on the day of euthanasia.

### Analysis of Tumor-infiltrating Lymphocytes (TIL) by Flow Cytometry

Tumors were collected on day 13, weighed, mechanically diced, and digested with liberase (2 mg/mL, 05401020001, Roche) and DNase I (50 μg/mL, 11284932001, Sigma-Aldrich) at 37°C for 30 minutes with rotation. Single-cell suspensions were obtained by filtering the digested tissues through a 45 μm strainer, after which erythrocytes were removed using 1x RBC lysis buffer (420301, Biolegend). To stain the cell surface markers of tumor-infiltrating lymphocytes (TILs), single-cell suspensions were blocked with anti-mouse CD16/32 (156603, BioLegend) for 10 minutes on ice, and then incubated with fluorochrome-labeled antibodies diluted with staining buffer (1:50, PBS, 2% FBS, 0.1% EDTA) for 30 minutes on ice in the dark. Dead cells were excluded using DAPI (1:1,000, BioLegend). After washing, cells were resuspended in 300-500 μL staining buffer for flow cytometry analysis.

Lymphoid cell phenotyping panel: CD45-APC750 (30-F11), CD3e-APC (145-2C11), CD4-BV650 (GK1.5), CD8-PercpCy5.5 (53-6.7), PD1-PE (RMP1-20), DAPI.

Myeloid cell phenotyping panel:CD45-APC750, I-A/I-E-APC (M5/114.15.2), CD11c-PE (N418), CD11b-PercpCy5.5 (M1/70), Ly6C-AF700 (HK1.4), F4/80-FITC (BM8), XCR1-BV650 (ZET).

All data were acquired using LSRII Analyzer and analyzed with Flow Jo v10.0.

### Cytokine array and enzyme-linked immunosorbent assay (ELISA)

Mouse serum was analyzed using multiplex immunoassays designed for mice (Mouse Cytokine Array / Chemokine Array 31-Plex (MD31) from Eve Technologies), with 8 replicates from each group. Heatmaps display relative cytokine expression values normalized to vehicle-treated samples. Serum HMGB1 was analyzed by ELISA (NOVUS, NBP2-62767) according to the manufacturer’s protocol.

### Immunohistochemistry and immunofluorescence

The mouse tumors were fixed in 4% paraformaldehyde overnight at 4°C, washed, and then stored in 70% ethanol until paraffin embedding. Paraffin sections (5μm) were hydrated for subsequent analysis.

For H&E staining, the hydrated slides were stained with hematoxylin and eosin. For IHC analysis, after hydration, the sections were subjected to antigen retrieval by incubating in citrate buffer (pH 6.0), Tris EDTA buffer (pH 9.0), or EDTA buffer (pH 8.0) at 121°C for 15 minutes. Endogenous peroxidase was blocked with 3% H2O2 in PBS, and non-specific binding was blocked with 2.5% normal serum for 1 hour at room temperature. The sections were then incubated with the respective primary antibodies overnight at 4°C. The primary antibodies used in IHC were cleaved caspase 3 (1:200, 9661, CST) and cleaved caspase 7 (1:200, 8438, CST). Following PBS washes, the sections were incubated with secondary antibodies (SignalStain Boost IHC Detection Reagent, 8114, CST), developed with DAB (SignalStain DAB Substrate,8059, CST), and counterstained with hematoxylin (VWR, 100504-658). At least four representative images of tumor sections from each group were acquired.

For immunofluorescence analysis, slides after antigen retrieval were incubated overnight at 4°C with the following primary antibodies: CD8 (1:100, 14-0808-82, eBioscience), CD4 (1:100, NBP1-19371, Novus Biologicals), FOXP3 (1:100, NB100-39002, Novus Biologicals), or F4/80 (1:100, NB600-404, Novus Biologicals). After washing with PBS, the tumor sections were incubated with Alexa Fluor 488/594/647 secondary antibodies (1:500; Thermo Fisher Scientific) for 1 hour at room temperature, followed by nuclei staining with DAPI for 20 minutes (1:30,000, 422801, Biolegend). The slides were mounted with ProLong™ Diamond Antifade Mountant (Thermo Fisher, P36970) and imaged using a Zeiss LSM780 confocal microscope with a ×60 objective. Consistent image exposure times and threshold settings were applied for all groups.

## Acknowledgement

This research was supported in part by the National Institute of Health (R01-CA250503 and R01-CA268518 to J.W.), the Cancer Prevention & Research Institute of Texas (CPRIT, RP220480 to J.W.), Howard Hughes Medical Institute (to M.C.W.), and Michael E. DeBakey, M.D., Professor in Pharmacology (to J.W.). We also appreciate all suggestions and advice from Drs. Ying Li, Douglas Green, Pascal Meier, Jeff Rosen, and Xiang Zhang.

## Data Access Statement

Research data supporting this publication are available upon request. The mass spectrometry raw data files for quantitative multiplexed proteomics have been deposited in the MassIVE dataset under accession number *MSV000092377*.

---

**Figure S1.**
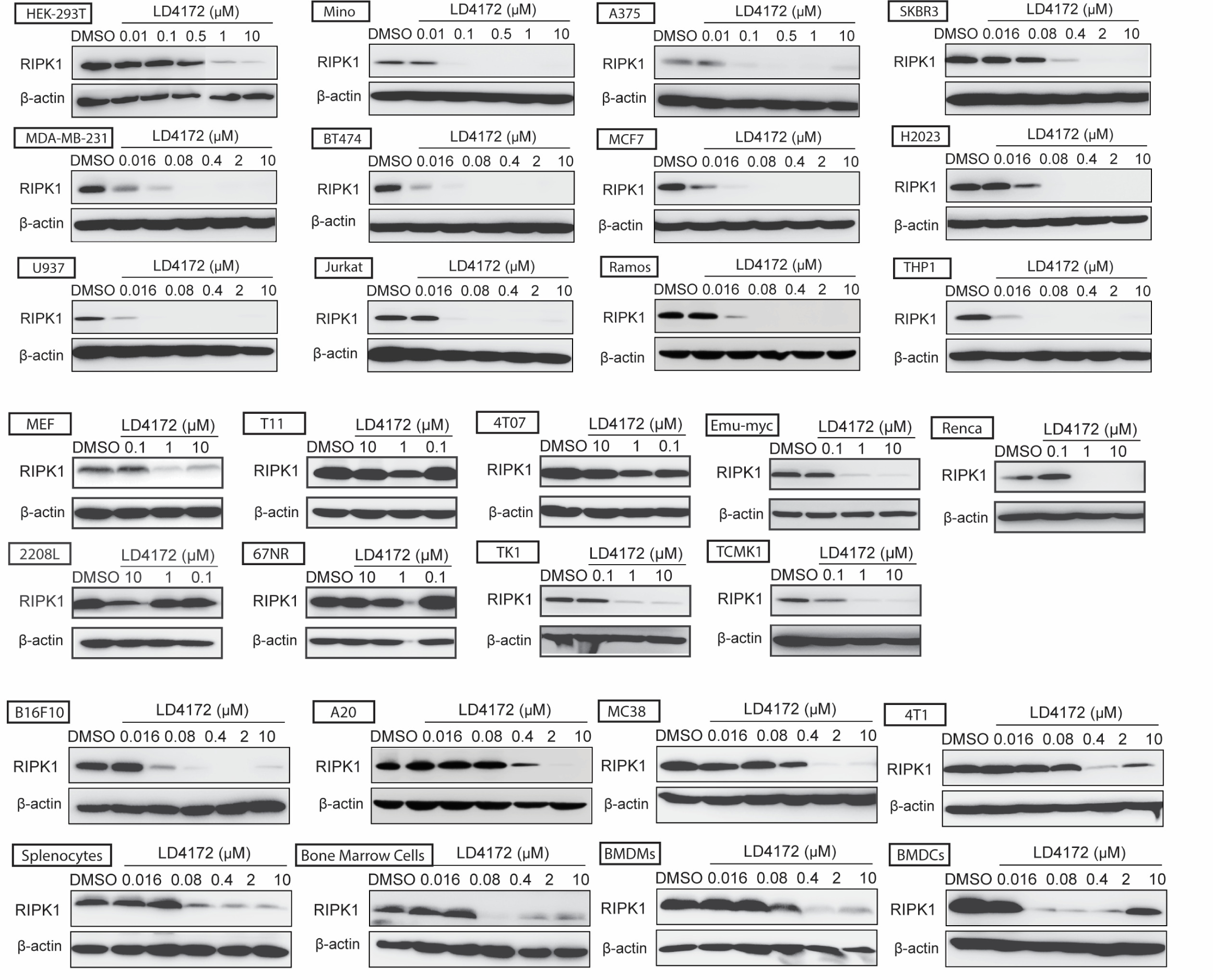
Degradation potency of LD4172 in a panel of cell lines. Representative Western blot assessing RIPK1 level in various cell lines treated with LD4172 at indicated concentrations for 24 h. The degradation potency of LD4172 in different cell lines was quantified using DC_50_ value and D_max_, which are listed in Fig. 2C.

**Figure S2.**
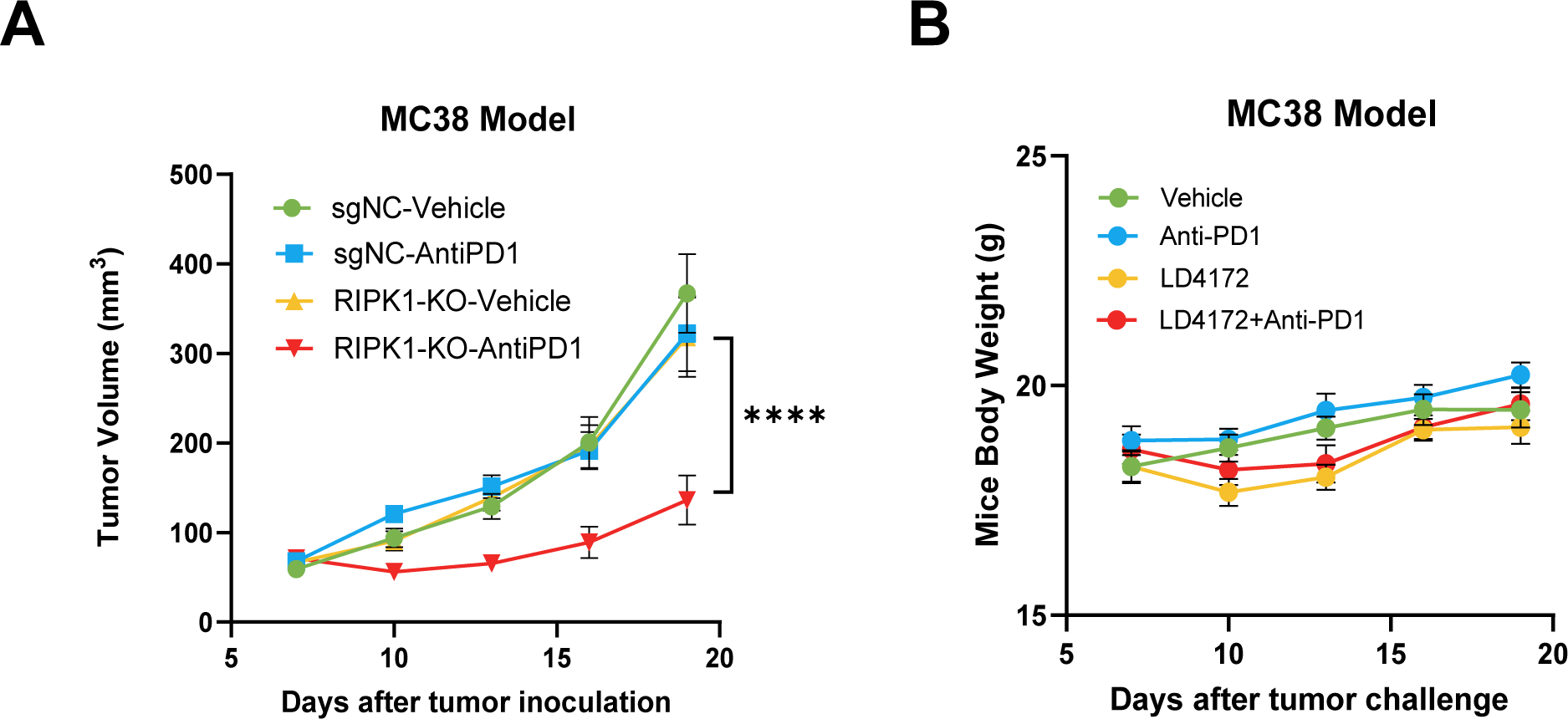
*In Vivo* efficacy of LD4172 in MC38 mouse model. **A,** Tumor growth curve of mice with MC38 tumors treated with LD4172 and/or anti-PD1 (n=18-36, 3 independent experiments). C57B6/J mice were subcutaneously inoculated with 3×10^5^ MC38 tumor cells. After seven days (tumor size ∼ 80 mm^3^), mice were treated every three days with anti-PD1 (100 µg per dose, i.p.); or daily with LD4172 (20 mg/kg, i.p.), or combination of LD4172 and anti-PD1 (same dose as their individual doses), or their corresponding vehicle control. **B,** Body weight curve of mice with indicated treatment.

**Figure S3.**
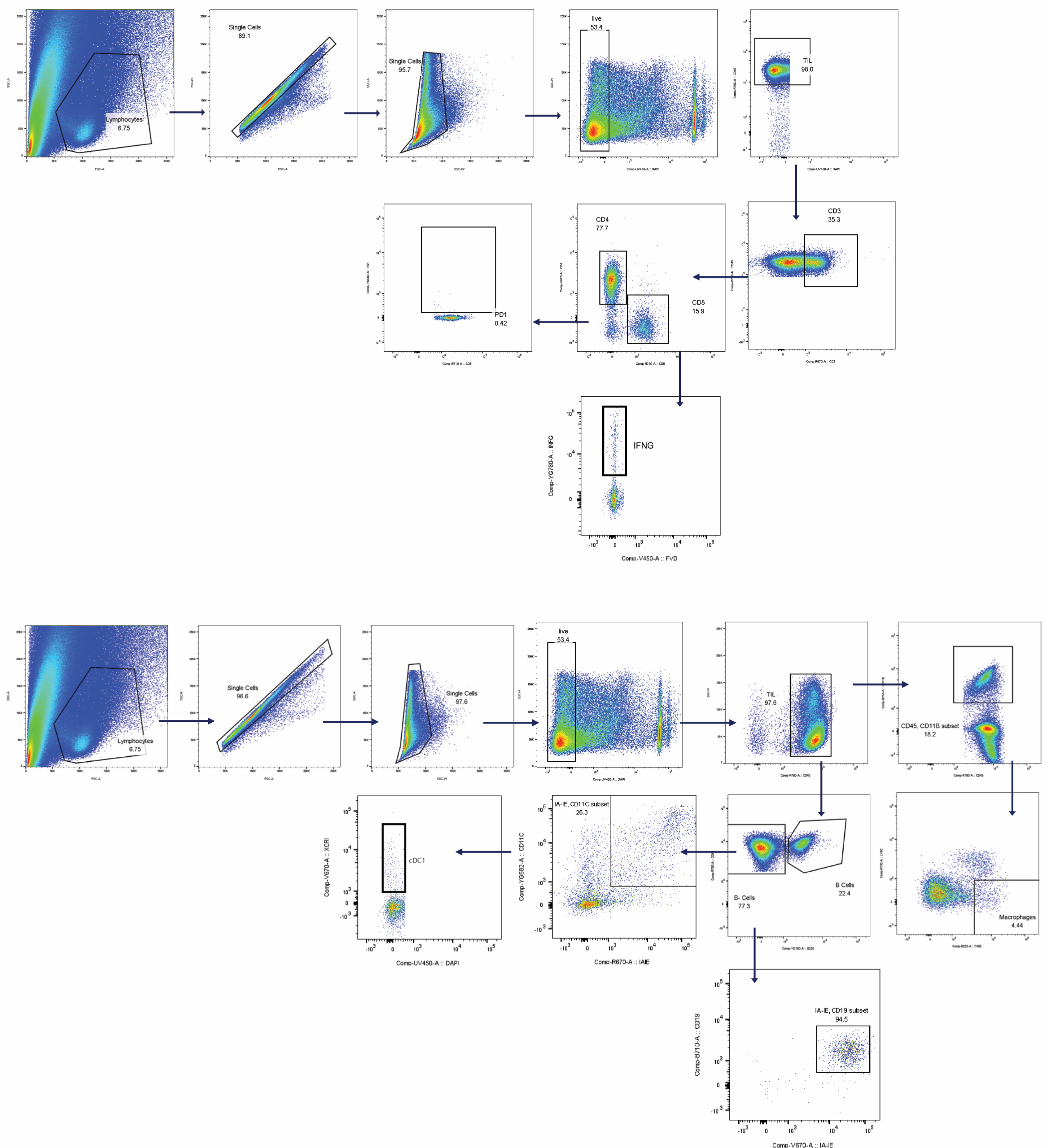
Representative plots showing the gating strategy for the data shown in Fig. 3.

**Figure S4.**
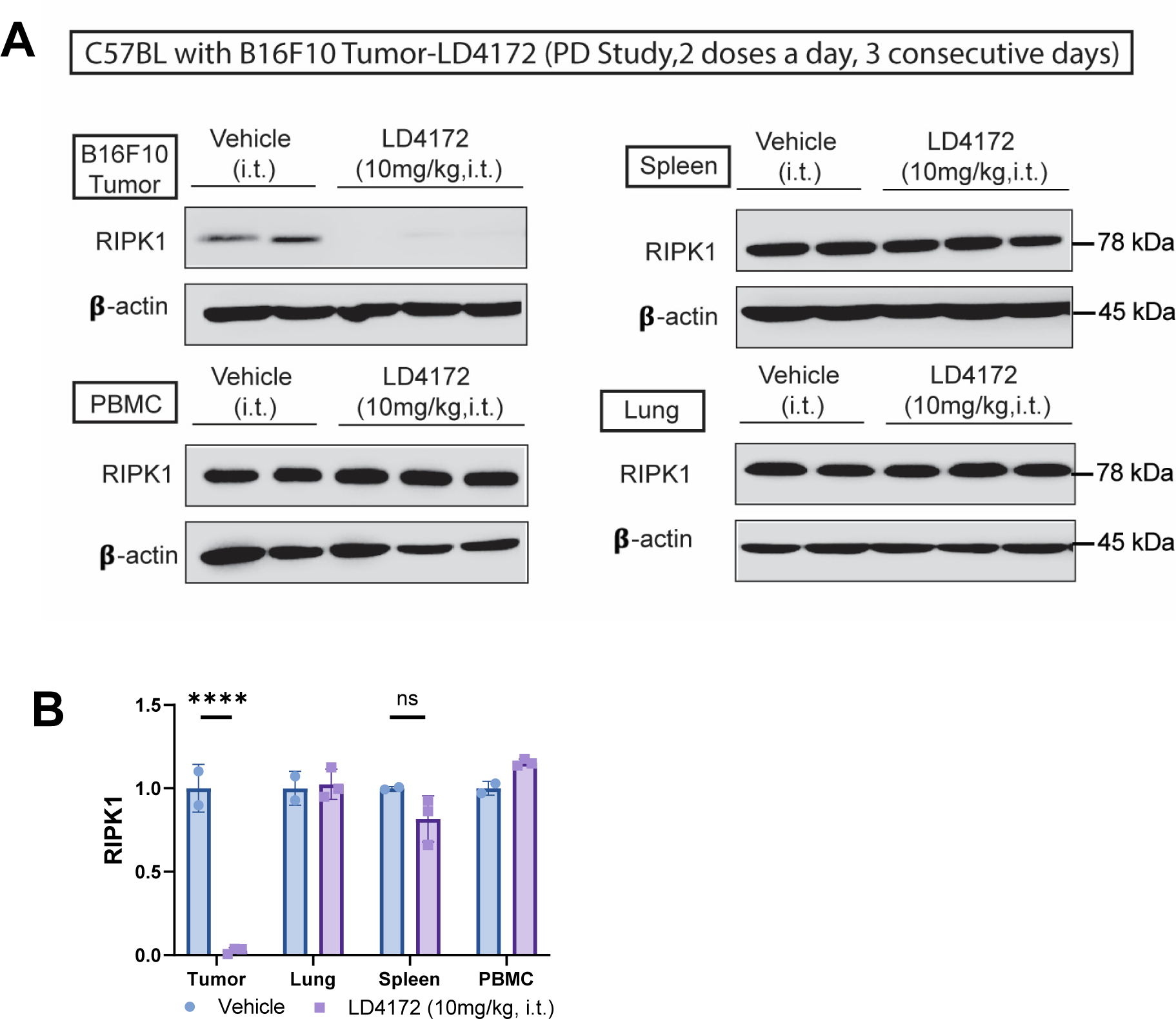
Pharmacodynamic (PD) properties of LD4172 with intratumoral administration. **A**, Representative immunoblots of RIPK1 expression in various tissues of C57BL/6J mice treated with LD4172 are shown. Mice with syngeneic B16F10 tumors received intratumoral injections of LD4172 (10mg/kg) twice daily for three days. Upon sacrifice, tissues were collected, and the levels of RIPK1 were quantified through Western blotting. **B**, Densitometric analysis of RIPK1 protein levels in different tissues is presented (n=2 or 3). Representative data from a minimum of two independent experiments are included. The data are presented as mean ± standard deviation (SD). Statistical significance is denoted as follows: * p<0.01; ** p<0.001; **** p<0.0001. ns indicates no statistical significance.

**Figure S5.**
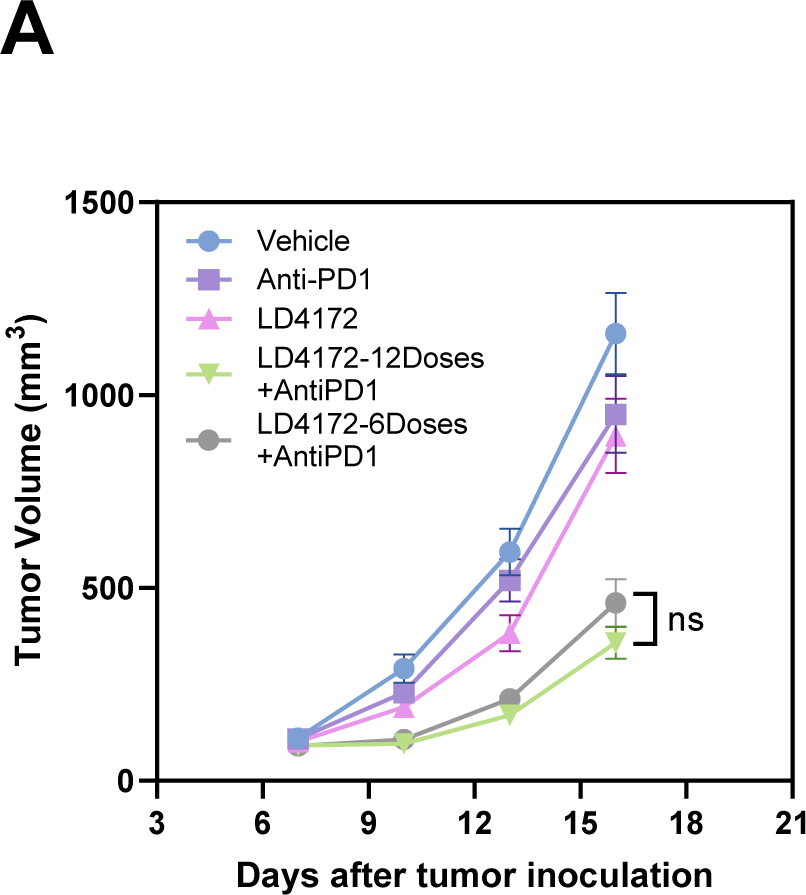
Tumor growth curve of mice with B16F10 tumors treated with reduced dosing frequency of LD4172. The administration of LD4172 was modified to a reduced dosage of 20 mg/kg every other day (n=8). The data are expressed as the mean ± SEM. * p<0.1; ** p<0.01; *** p<0.001; **** p<0.0001. ns, no statistical significance.

## References

1. Korman, A. J., Garrett-Thomson, S. C. & Lonberg, N. The foundations of immune checkpoint blockade and the ipilimumab approval decennial. Nat Rev Drug Discov 21, 509–528 (2022).

2. Vesely, M. D., Zhang, T. & Chen, L. Resistance Mechanisms to Anti-PD Cancer Immunotherapy. Annual Review of Immunology 40, 45–74 (2022).

3. Upadhaya, S., Neftelinov, S. T., Hodge, J. & Campbell, J. Challenges and opportunities in the PD1/PDL1 inhibitor clinical trial landscape. Nature Reviews Drug Discovery 21, 482–483 (2022).

4. Mifflin, L., Ofengeim, D. & Yuan, J. Receptor-interacting protein kinase 1 (RIPK1) as a therapeutic target. Nat Rev Drug Discov 19, 553–571 (2020).

5. Cucolo, L. et al. The interferon-stimulated gene RIPK1 regulates cancer cell intrinsic and extrinsic resistance to immune checkpoint blockade. Immunity 55, 671–685.e10 (2022).

6. Hou, J. et al. Integrating genome-wide CRISPR immune screen with multi-omic clinical data reveals distinct classes of tumor intrinsic immune regulators. J Immunother Cancer 9, e001819 (2021).

7. Manguso, R. T. et al. In vivo CRISPR screening identifies Ptpn2 as a cancer immunotherapy target. Nature 547, 413–418 (2017).

8. Shi, K., Zhang, J., Zhou, E., Wang, J. & Wang, Y. Small-Molecule Receptor-Interacting Protein 1 (RIP1) Inhibitors as Therapeutic Agents for Multifaceted Diseases: Current Medicinal Chemistry Insights and Emerging Opportunities. J. Med. Chem. 65, 14971–14999 (2022).

9. Burslem, G. M. & Crews, C. M. Small-Molecule Modulation of Protein Homeostasis. Chemical reviews 117, 11269–11301 (2017).

10. Li, Y. et al. Identification of 5-(2,3-Dihydro-1 *H*-indol-5-yl)-7 *H*-pyrrolo[2,3-*d*] pyrimidin-4-amine Derivatives as a New Class of Receptor-Interacting Protein Kinase 1 (RIPK1) Inhibitors, Which Showed Potent Activity in a Tumor Metastasis Model. J. Med. Chem. 61, 11398–11414 (2018).

11. Yoshikawa, M. et al. Discovery of 7-Oxo-2,4,5,7-tetrahydro-6 H-pyrazolo[3,4-c] pyridine Derivatives as Potent, Orally Available, and Brain-Penetrating Receptor Interacting Protein 1 (RIP1) Kinase Inhibitors: Analysis of Structure–Kinetic Relationships. J. Med. Chem. 61, 2384–2409 (2018).

12. Robers, M. B. et al. Target engagement and drug residence time can be observed in living cells with BRET. Nat Commun 6, 10091 (2015).

13. Guo, W.-H. et al. Enhancing intracellular accumulation and target engagement of PROTACs with reversible covalent chemistry. Nat Commun 11, 4268 (2020).

14. Huang, H. T. et al. A Chemoproteomic Approach to Query the Degradable Kinome Using a Multi-kinase Degrader. Cell chemical biology 25, 88–99 e6(2018).

15. Dillon, C. P. et al. RIPK1 Blocks Early Postnatal Lethality Mediated by Caspase-8 and RIPK3. Cell 157, 1189–1202 (2014).

